# Influence of SNF1 complex on growth, glucose metabolism and mitochondrial respiration of *Saccharomyces cerevisiae*

**DOI:** 10.1101/287961

**Authors:** Cecilia Martinez-Ortiz, Andres Carrillo-Garmendia, Blanca Flor Correa-Romero, Melina Canizal-García, Juan Carlos González-Hernández, Carlos Regalado-Gonzalez, Ivanna Karina Olivares-Marin, Luis Alberto Madrigal-Perez

## Abstract

The switch of mitochondrial respiration to fermentation as the main pathway to produce ATP through the increase of glycolytic flux is known as the Crabtree effect. The elucidation of the molecular mechanism of the Crabtree effect may have important applications in ethanol production and lay the groundwork for the Warburg effect, which is essential in the molecular etiology of cancer. A key piece in this mechanism could be Snf1p, which is a protein that participates in the nutritional response that includes glucose metabolism. Thus, this work aimed to recognize the role of the SNF1 complex on the glycolytic flux and mitochondrial respiration, to gain insights about its relationship with the Crabtree effect. Herein, we found that in *Saccharomyces cerevisiae* cells grown at 1% glucose, mutation of *SNF1* gene decreased glycolytic flux, increased NAD(P)H, enhanced *HXK2* gene transcription, and decreased mitochondrial respiration. Meanwhile, the same mutation increased the mitochondrial respiration of cells grown at 10% glucose. Moreover, *SNF4* gene deletion increased respiration and growth at 1% of glucose. In the case of the *GAL83* gene, we did not detect any change in mitochondrial respiration or growth. Altogether, these findings indicate that *SNF1* is vital to switch from mitochondrial respiration to fermentation.

## Introduction

One of the most relevant metabolic capacities of *Saccharomyces cerevisiae* is the ability to grow under aerobic and anoxic conditions; this yeast has evolved to use complex mechanisms to obtain energy in both circumstances switching the metabolism from respiration to mainly fermentation. The amount of fermentable sugars in the medium also promotes the preference in energy production of *S. cerevisiae*. Fermentable sugars concentration >0.8 mM favors energy generation through fermentation, while lower levels stimulate mitochondrial respiration (De Deken, 1966, Hagman, *et al.*, 2014). Interestingly, when glucose availability is enough, the yeast prefers the fermentative metabolism even in the presence of oxygen, and represses mitochondrial respiration, this phenotype is known as the Crabtree effect (Moriya & Johnston, 2004). Nonetheless, the metabolic regulation that governs these changes is not entirely understood. The increase of fermentable sugars availability inactivates the sucrose non-fermenting protein-1 (Snf1p) suggesting that this protein plays a vital role in establishing fermentation as the primary source of ATP production (Kayikci & Nielsen, 2015) (Celenza & Carlson, 1984).

The Snf1p protein is orthologous to the mammalian AMP-kinase (AMPK) (Hedbacker & Carlson, 2008). Snf1p is a catabolic regulator that could be activated by increasing the ADP/ATP ratio (Mayer, *et al.*, 2011), or through phosphorylation by the kinases Sak1p, Elm3p, and Tos3p (Hong, *et al.*, 2003, Kim, *et al.*, 2005, Liu, *et al.*, 2011). Snf1p activation has been linked to several catabolic pathways regulation such as mitochondrial respiration (Wright & Poyton, 1990), fatty acid metabolism (Zhang, *et al.*, 2013), and glycolysis (Nicastro, *et al.*, 2015). However, modulation of Snf1p is intimately related to the glycolysis function. In this regard, the mutation of *SNF1* gene has profound effects on glucose metabolism. For example, the *snf1*∆ mutant strain is unable of growing in media with low levels of glucose, but at high levels, the growth is similar to the wild type strains (Carlson, *et al.*, 1981). Additionally, the glycolytic flux is also impaired by deletion of the *SNF1* gene (Nicastro, *et al.*, 2015). A key player in the phenotype showed by the *snf1*∆ mutant is Mig1p, which is phosphorylated and inhibited by Snf1p and is responsible for regulating several proteins including the hexose transporters *HXT1* and *HXT3*. Accordingly, the *MIG1* mutation partially relieves the glucose repression (Treitel, *et al.*, 1998). Interestingly, the *snf1*∆ mutant strain is more susceptible to the toxicity induced by deoxyglucose (glucose analog that inhibits glycolysis), and the overexpression of the Hxt1p and Hxt3p release the mutant strain from this toxicity (O’Donnell, *et al.*, 2015). One of the most accepted theories about the Crabtree effect is that it originates by the hexose transporters modulated by the glycolytic flux (Huberts, *et al.*, 2012).

Thereby the glucose repression might be originated by the down-regulation of specific hexose transporters through the Snf1p/Mig1p pathway. Hence, our work aimed to determine whether deletion of the *SNF1* gene in *S. cerevisiae* impairs the glycolytic flow and how it affects the switch of mitochondrial respiration to fermentation due to glucose repression.

## Material and methods

### Strains

The experiments were carried out in the genetic background of *S. cerevisiae* BY4742 (*MaT*α, *his3*Δ*1, leu2*Δ*0, lys2*Δ*0, ura3*Δ*0*) and its mutant in the genes *SNF1* (*snf1*Δ, *MAT*α, *his3*Δ*1, leu2*Δ*0, lys2*Δ*0, ura3*Δ*0, YDR477w: kanMX4*), *SNF4* (*snf4*Δ, *MAT*α, *his3*Δ*1, leu2*Δ*0, lys2*Δ*0, ura3*Δ*0, YGL115w: kanMX4*), *GAL83* (*gal83*Δ, *MAT*α, *his3*Δ*1, leu2*Δ*0, lys2*Δ*0, ura3*Δ*0, YER027c: kanMX4*) acquired from the EUROSCARF program (University of Frankfurt, Germany). Strains were maintained in YPD medium (1% yeast extract, 2% casein peptone and 2% glucose). The mutant strains were supplemented with the antibiotic geneticin (200 mg/mL) (G-418, Sigma-Aldrich, St. Louis, MO, USA).

### Experimental design

To verify that Snf1p is mainly regulated by the concentration of carbon sources and not by nitrogen sources, we used a 3^4^ full factorial randomized design. The following factors were evaluated: type of carbon source (levels: glucose, sucrose and galactose), type of nitrogen source (levels: proline, glutamate, and ammonium sulfate), carbon concentration (levels (w/v): 0.01%, 2%, and 10%), and nitrogen concentration (levels (w/v): 0.01%, 0.5% and 5%). A total of 81 nutritional conditions were assayed three to five times with two technical replicates. The same experimental design was performed using the BY4742 and *snf1*Δ strains separately. The response variable was the percentages of growth (specific growth rate) of the mutant *snf1*Δ relative to the growth of WT multiplied by 100. Cultures were grown on honeycomb plates (Growth Curves, Piscataway, NJ, USA) with 145 μL of medium per well. Each well was inoculated with 5 μL of an overnight *S. cerevisiae* BY4742 or *snf1*Δ culture, grown in YPD medium at 30 °C in an orbital shaker (MaxQ 6,000, Thermo Scientific, Waltham, MA, USA) at 250 rpm. Samples were incubated in SC medium at 30 °C for 48 h using a Bioscreen (model C MBR, Growth Curves) programmed with continuous shaking at medium speed and readings at 600 nm, every 30 min. A dendogram was constructed using the average linkage clustering method and the Euclidean distance measurement method using the program heatmapper (Babicki, *et al.*, 2016).

### Plate serial dilutions

Each strain was grown in 3 mL of YPD medium supplemented with 2% glucose in 10 mL test tubes inoculated with three single colonies from YPD agar plates. The cells were kept in the incubator with constant shaking (200 rpm) for 18 hours at 30 °C. The inoculum optical density at 600 nm (OD_600_) was adjusted to a final value of ~0.3 for serial dilutions. Then, 3 μL of 10-fold serial dilutions were plated on YPD medium supplemented with two concentrations of glucose (1 and 0.1%). Plates were incubated at 30 °C for 48 h.

### Extracellular acidification rate

A 3 mL YPD medium was inoculated with a fresh colony grown on the YPD agar plates and cultured at 30 °C and 200 rpm overnight. The pre-culture was used to inoculate 250 mL flasks with 50 mL of YPD medium supplemented with 1 or 10% glucose. The cells were harvested at mid-log phase (OD_600_~0.5) at 9,000 *g* for 5 minutes, and three washes were performed with sterile water, and the cell pellet was resuspended in 2 mL of sterile water. Then, cells were added to 25 mL of distilled water to obtain a final concentration of about 120 mg cells/mL. The extracellular acidification rate experiment was initiated by adding 1 mL of 1 M glucose, and recording the pH in the Metrohm 902 Titrando equipment (Metrohm, FL, USA), fitted with the Tiamo software version 2.3 for 5 minutes. Data were analyzed with the GraphPad Prism 6.00 for Macintosh (GraphPad Software, La Jolla California, USA).

### Quantification of NADH/NAD^+^

Cells were cultured in 10 mL of YPD medium supplemented with 1% or 10% (w/v) glucose at 30 °C with constant agitation (200 rpm), when cells reached the mid-log phase (OD_600_~0.5) 35 mL of methanol was added. Then, the solution was centrifuged at 4,000 *g* for 1 minute at −20 °C, immediately resuspended in 0.25 mL of 0.1 N NaOH (NADH) or 0.1 N HCl (NAD^+^), boiled for 1 minute and centrifuged at 5,000 *g* for 5 minutes to separate and discard the cellular debris. Concentrations of NADH and NAD^+^ were determined according to Bernofsky and Swan (1973).

### RT-qPCR

The strains were grown in 3 mL of YPD medium supplemented with 1 or 10% glucose until the mid-log phase (OD_600_~0.5) was reached. Subsequently, the cells were harvested to remove the culture medium, centrifuging at 13,000 *g* for 3 minutes. Then, 0.75 mL of TRIzol (Ambion, Life Technologies, Foster City, CA, USA) was added (the cell pellet was not washed with TRIzol to avoid lysis and release of nucleases), homogenized at 3,000 rpm for 15 seconds, and immediately placed in ice. The suspension was centrifuged at 12,000 *g* for 10 minutes at 4 °C, and the supernatant was transferred to a new microtube, followed by incubation for 5 minutes at room temperature, to allow complete dissociation of the nucleoprotein complexes. Later, 0.2 mL of cold chloroform was added per mL of TRIzol, hand shaken vigorously for 15 seconds and incubated for 2-3 minutes at room temperature, followed by centrifugation at 12,000 *g* for 15 minutes at 4 °C. The aqueous phase was immediately transferred to a sterile microtube where 0.5 mL of 100% isopropanol was added and incubated for 10 minutes, before centrifuging at 12,000 *g* for 10 minutes at 4 °C, and the supernatant was removed. The pellet was washed with 1 mL of 75% ethanol, homogenized and then centrifuged at 7,500 *g* for 5 minutes at 4 °C. The pellet was allowed to dry, and then 50 μL diethylpyrocarbonate water (DEPC), and recombinant DNAase I, RNAse free (Roche Diagnostics GmbH, Manheim, Germany) were added. The integrity of the RNA was verified with an electrophoretic analysis, as well as the concentration and quality of the RNA using a Nanodrop 2000 (Thermo-Scientific, Whaltham, MA, USA).

To carry out the qPCR, the complementary DNA synthesis (cDNA) was performed. We used 2.5 μg of total RNA and the RevertAid H Minus first chain cDNA synthesis kit (Thermo-Scientific). First, 1 μL of random hexamer primer (200 ng), 12 μL of DEPC water and 1 μL of dNTP (10 mM) were added. It was mixed and heated at 65 °C for 5 minutes, allowed to cool on ice, added with 4 μL of 5x buffer, and 2 μL of DTT (0.1 M), mixed and heated at 37 °C for 2 minutes. Then 1 μL of RevertAid H Minus M-MuLV RT was added, allowed to incubate at room temperature for 10 minutes. Then, it was heated at 37 °C for 50 minutes and finally heated at 70 °C for 15 minutes. Once the cDNA was synthesized, the RT-qPCR was continued. Each reaction had a final volume of 20 μL, 2 μL of cDNA, 10 μL of SYBR Select Master Mix (AppliedBiosystems), 0.8 μL of each primer and 6.4 μL of DEPC water were added. Subsequently, they were placed in the Rotor-Gene Q (Qiagen, Venlo, The Netherlands).

The genes that were evaluated at the transcriptional level were *HXK2* and *PFK1*, using as a reference gene *UBC6* (ubiquitin-conjugation-6). Conditions used to carry out the qPCR were those depicted by the protocol of Madrigal-Perez, *et al.* (2015) which were: beginning with the activation of UDG at 50 °C for 2 minutes, followed by the activation of AmpliTaq-DNA at 95 °C for 2 minutes with 35 cycles of denaturation at 94 °C for 15 seconds. The alignment temperature of oligos *HXK2* and *UBC6* was 51 °C, oligo *PFK1* was 54 °C for 30 seconds, and the extension temperature of 72 °C for 1 minute, followed by 1 cycle at 72 °C for 7 minutes.

### *Determination of* in situ *mitochondrial respiration*

The mitochondrial respiration was performed according to Tello-Padilla, *et al.* (2017). Briefly, the strains were cultured in 50 mL of YPD medium supplemented with (w/v) 1% or 10% glucose at 30 ºC and 200 rpm until the mid-log phase (OD_600_~0.5) was reached. Cells were harvested at 5,000 *g* for 5 minutes, and three washes were performed with deionized water and resuspended in a 1:1 ratio (w/v). Oxygen consumption was analyzed polarographically with a Clark type electrode connected to a YSI5300A monitor (Yellow Springs, OH, USA) and a computer for data acquisition. In the polarograph chamber 125 mg of cells (wet weight), 5 mL of MES-TEA buffer (10 mM morphoethanolsulfonic acid, pH 6.0 with triethanolamine) and glucose (10 mM) were added; the cells were maintained with constant agitation (basal respiration). Subsequently, the uncoupled state (maximum respiration) was stimulated with the addition of 0.015 mM carbonylcyanide-*p*-trichlorophenylhydrazone (CCCP) for 3 minutes. Then, electron transport chain (ETC) inhibitors were added: thenoyltrifluoroacetone (TTFA, 1 mM), antimycin A (AA) and 0.75 mM KCN (non-mitochondrial respiration), each inhibitor was left for 3 minutes. The results were analyzed using the statistical package GraphPad Prism 6.00 for Macintosh (GraphPad Software).

## Results

### *Effect of the chemical nature and concentration of carbon and nitrogen in growth of the mutant* snf1Δ

To verify that Snf1p is mainly regulated by the concentration of carbon sources and not by nitrogen sources, we made an experimental design with three concentrations of nitrogen and carbon sources using three different carbon sources (galactose, glucose, and sucrose) and three nitrogen sources (proline, ammonium, and glutamate). We found that the main factor affecting the growth of the mutant *snf1*Δ was the carbon source concentration (Fig. 1). However, the nitrogen source and its concentration do not show any effect upon the growth of the mutant *snf1*Δ (Fig. 1). Growth decrease is evident at ≥0.5% glucose (Fig. 1). These data corroborate the critical role of *SNF1* gene in the growth of *S. cerevisiae* under low concentration of carbon sources.

**Figure 1.**
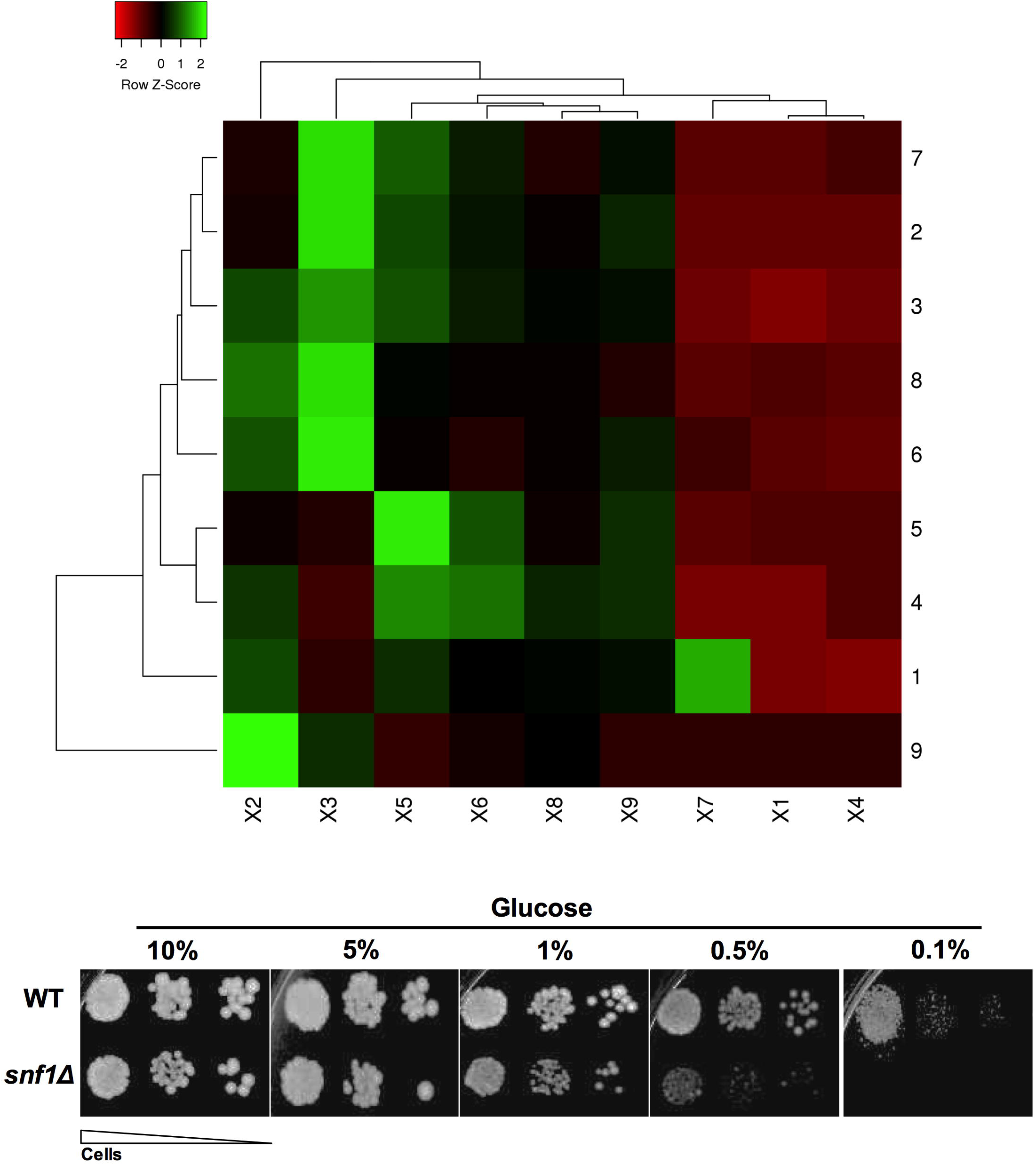
Influence of chemical nature and concentration of carbon and nitrogen sources on growth of the mutant *snf1*Δ. Data in the heat map shows the cluster membership of the mutant *snf1*Δ growth relative to the WT strain times 100. Glucose supplementation at 0.01%, 2%, and 10% are represented by X1, X2, and X3. Galactose supplementation at 0.01%, 2%, and 10% are represented by X4, X5, and X6. Sucrose supplementation at 0.01%, 2%, and 10% are represented by X7, X8 and X9. Proline supplementation at 0.01%, 0.5%, and 5% are represented by 1, 2, and 3. Glutamate supplementation at 0.01%, 0.5%, and 5% are represented by 4, 5, and 6. Ammonium sulfate supplementation at 0.01%, 0.5%, and 5% are represented by 7, 8, and 9. The average linkage clustering method was used, while Euclidean distance measurement method was used. The results represent mean values from 3-5 independent experiments, which includes mean values of 3 technical repetitions.

### *Impact of the* SNF1 *mutation in the glycolytic flux*

To obtain the first insights about how the *SNF1* gene affects glucose metabolism, we decided to measure the extracellular acidification rate as an indicator of the glycolytic flux. Two glucose concentrations were used (1% and 10%) since concentrations below 0.5% glucose the mutant *snf1*Δ showed a negligible growth. Interestingly, at 1% glucose the mutant *snf1*Δ showed a lower acidification rate than the WT strain (Fig. 2), while at 10% glucose no differences were revealed (Fig. 2). These data proved that the *SNF1* gene is vital to sustain the glycolytic flux in a low glucose concentration.

**Figure 2.**
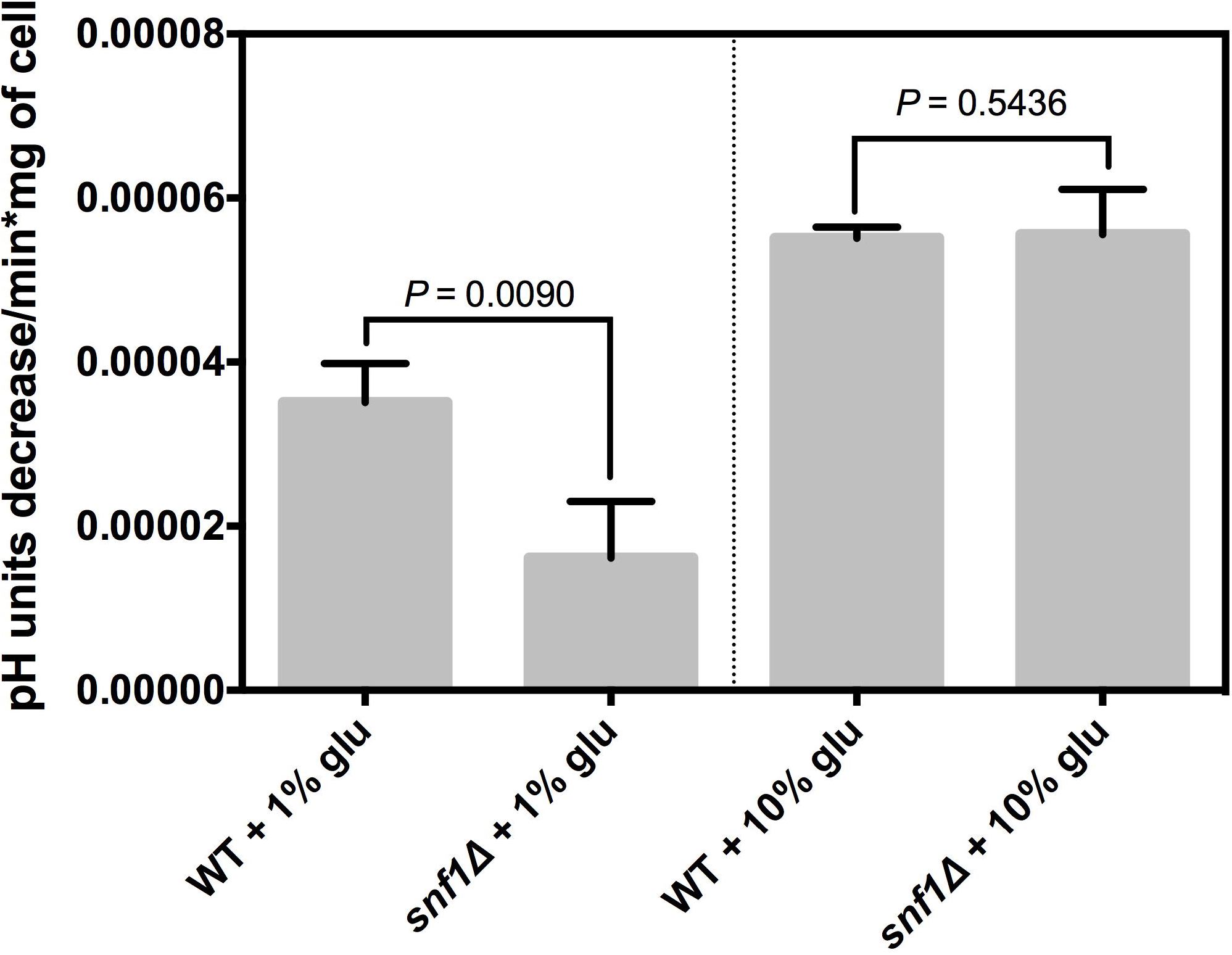
Effect of the *SNF1* deletion in the extracellular acidification rate. Media acidification was used as an indirect indicator of glucose consumption at 1% and 10% glucose. The results represent mean values ± SEM from 3-4 independent experiments. Statistical analyses were performed using two-tailed paired Student *t-*test.

### *Influence of* SNF1 *mutation on NADH/NAD^+^ and NAD(P)H ratios*

A useful indicator of how glycolysis is operating NADH/NAD^+^ ratio since it is a key regulator of this pathway. This parameter was measured, to discard whether the decrease of the glycolytic flux is due to a down-regulation of the glycolysis pathway. No differences were found either at 1% or 10% of glucose in the NADH/NAD^+^ ratio (Fig. 3). Another possibility is that glucose-6-phosphate has been metabolized by the pentose phosphate pathway, to prove this idea the NAD(P)H was measured, since we did not find any difference in the NADH/NAD^+^ ratio. Therefore glycolysis changes could be attributed to NADPH variations. At 1% glucose, we found higher NAD(P)H concentration in the mutant *snf1*Δ than in the WT strain (Fig. 3). However, no differences were observed at 10% glucose. These data suggest that a decrease of the glycolytic flux of the mutant *snf1*Δ could be due to a bypass of glycolysis to the pentose phosphate pathway at 1% glucose.

**Figure 3.**
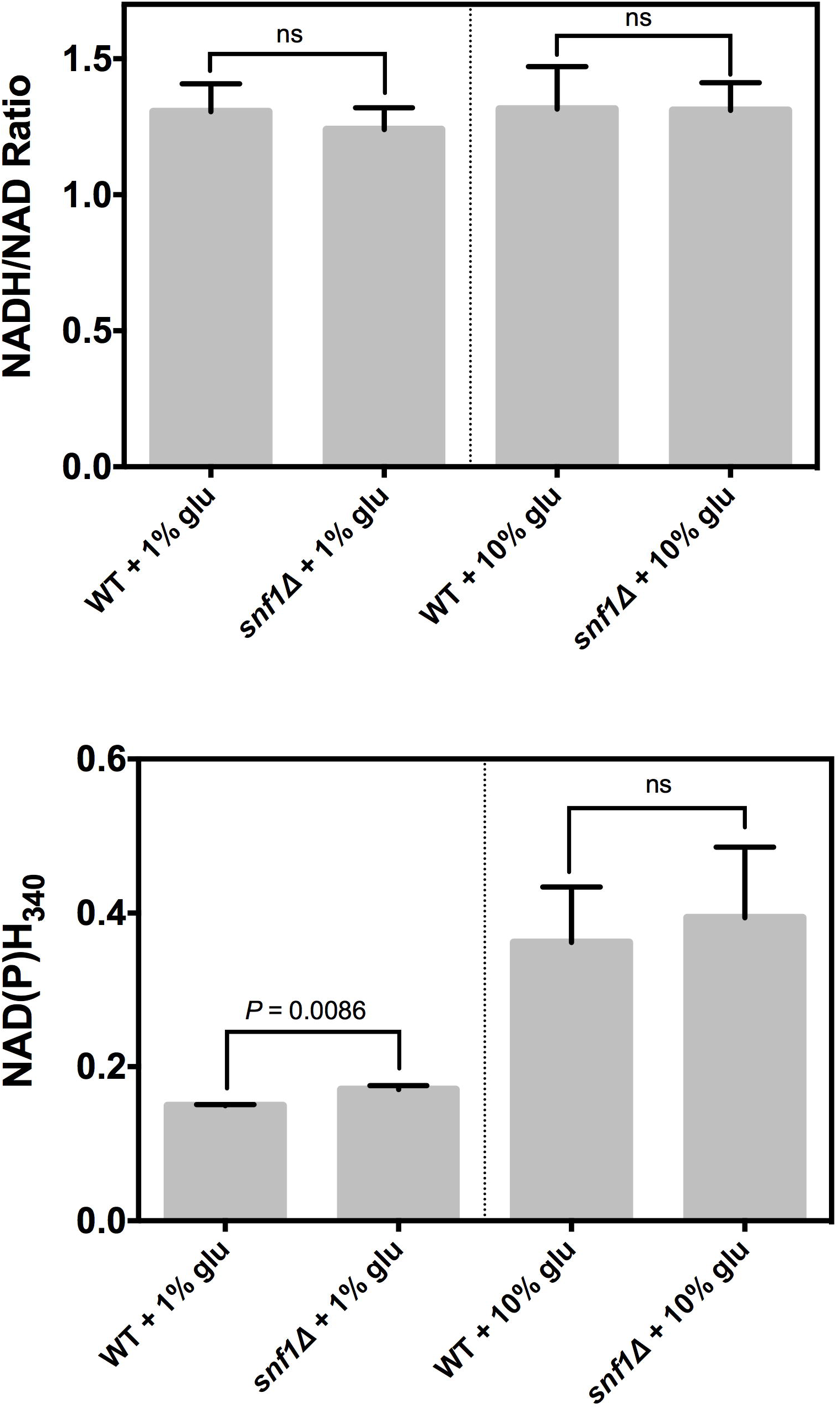
Variation of the NADH/NAD^+^ ratio and NAD(P)H in the mutant *snf1*Δ. NADH/NAD^+^ ratio measurement was performed using a cycling assay. NAD(P)H was calculated by fluorescence at 340 nm. The results represent mean values ± SEM from 3 independent experiments, which include mean values of 3 technical repetitions. Statistical analyses were performed using two-tailed paired Student *t-*test; ns, not significant.

### SNF1 *deletion affects the transcription of the* HXK2 *and* PFK1 *genes*

Mutation of the *SNF1* gene could affect the transcription of *HXK2* and *PFK1*, two critical genes of the glycolysis pathway that in turn possibly results in the glycolytic flux impairment. Regarding the *HXK2* gene, it was found that transcript levels in the mutant *snf1*Δ were higher than in the WT strain at 1% glucose; while no differences were found at 10% glucose (Fig. 4). *PFK1* transcript levels did not change under any condition (Fig. 4). These data indicate that higher transcription levels of the *HXK2* gene could be related to the decrease of glycolytic flux seen in the mutant *snf1*Δ at 1% glucose.

**Figure 4.**
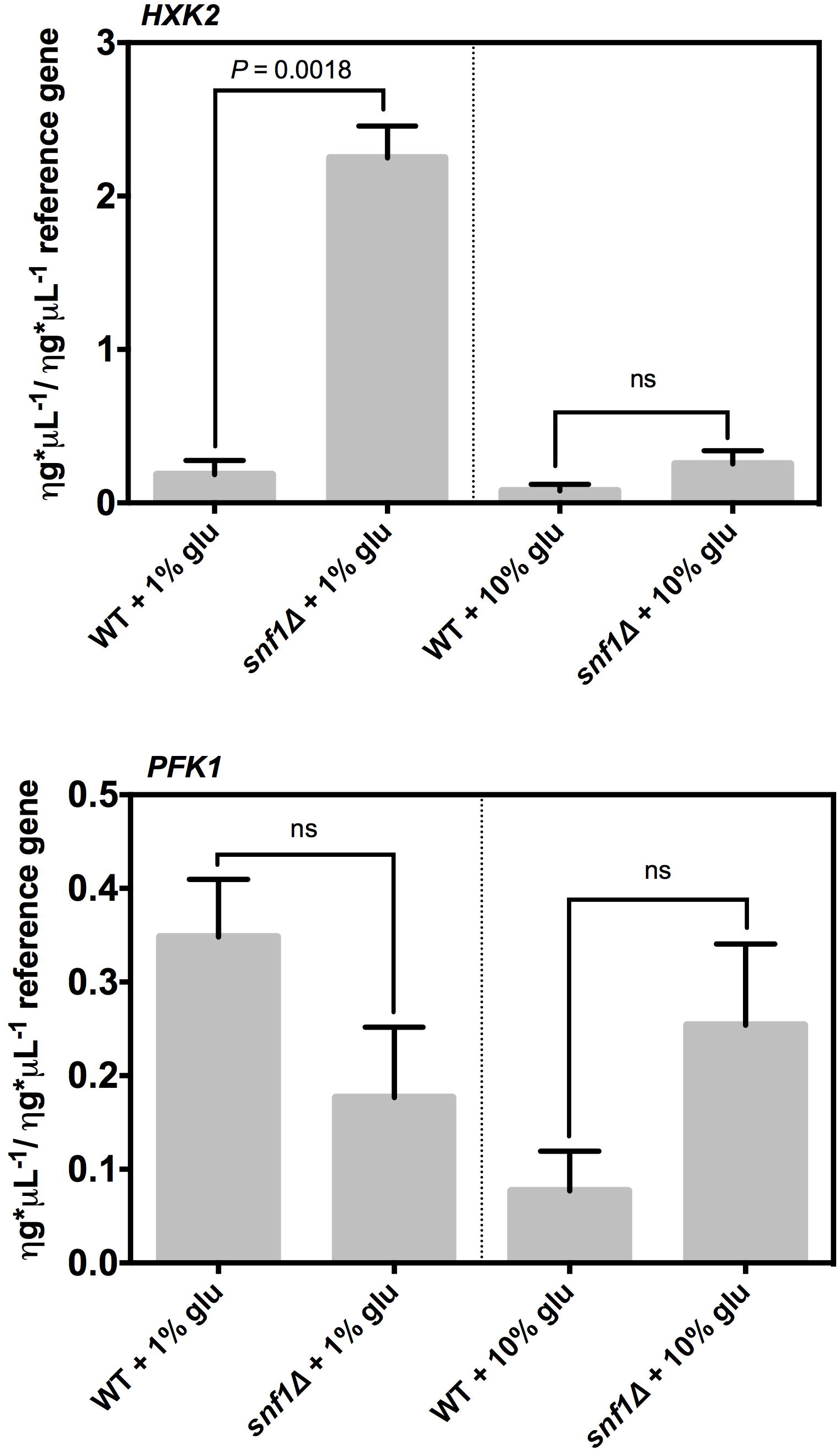
Transcription of genes *HXK2* and *PFK1* by deletion of the *SNF1* gene. Real-time PCR (qPCR) was performed to measure mRNA levels of *HXK2* and *PFK1* genes and *UBC6* as a reference gene. The results represent mean values ± SEM from 3 independent experiments, which include mean values of two technical repetitions. Statistical analyses were performed using two-tailed paired Student *t-*test; ns, no significance.

### *Effect of* SNF1 *mutation in the mitochondrial respiration*

The glycolytic flux is tightly coupled with mitochondrial respiration in *S. cerevisiae.* For example, the Crabtree effect is characterized by a decrease of the mitochondrial respiration and an increase of the glycolytic flux. To further understand the role of the gene *SNF1* in this phenotype, the mitochondrial respiration was measured. Interestingly, basal respiration and the maximal respiratory capacity were reduced in the mutant *snf1*Δ at 1% glucose in comparison with the WT strain (Fig. 5). Surprisingly, at 10% glucose the opposite phenotype was seen, the basal respiration and the maximal respiratory capacity were higher in the mutant *snf1*Δ than in the WT strain (Fig. 5). These data indicate that mitochondrial respiration is affected by *SNF1* mutation in a glucose-dependent manner.

**Figure 5.**
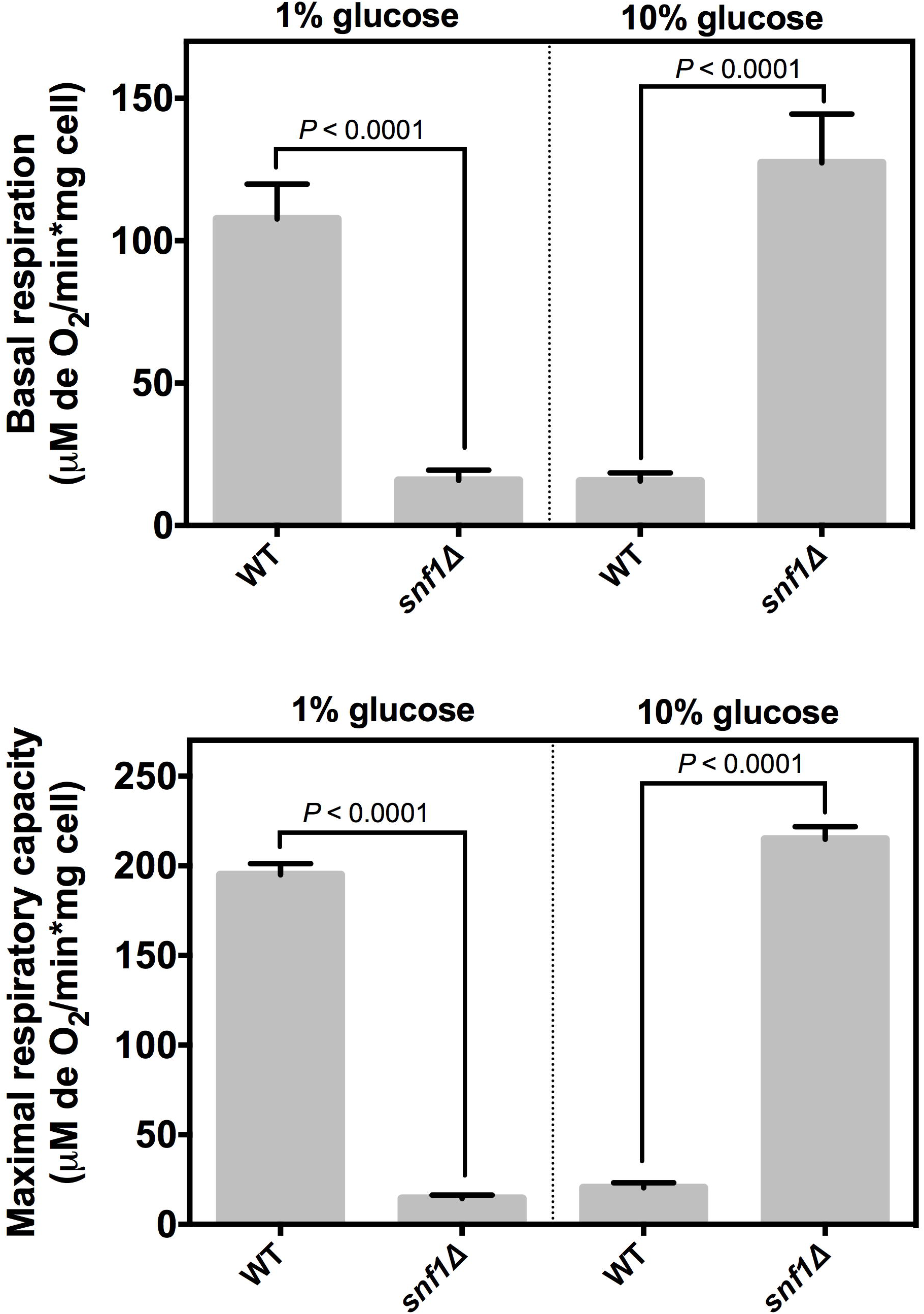
Influence of *SNF1* deletion in the mitochondrial respiration. The oxygen consumption was measured at basal state and maximal respiratory capacity in exponential phase. The results represent mean values ± SEM from 3-5 independent experiments, which includes mean values of 3 technical repetitions. Statistical analyses were performed using two-tailed unpaired Student *t-*test; ns, not significant.

### Influence of the SNF1 complex in the mitochondrial respiration

Snf1p is part of the SNF1 complex, which structurally is a heterotrimer consisting of the catalytic subunit Snf1p (α), the regulatory subunit Snf4p (γ) and three alternative subunits Gal83p, Sip1p and Sip2p (β). It has been shown that Snf4p binds to the regulatory domain in the Snf1p protein preventing the autoinhibition of the kinase activity of Snf1p under low glucose concentrations (Jiang & Carlson, 1996). The regulatory subunits Snf4p (γ) and Gal83p (β) have crucial participation in the activation and correct function of Snf1p. To understand the participation of the SNF1 complex in the mutants *snf4*Δ and *gal83*Δ the mitochondrial respiration was measured. Snf4p prevents the inhibition of Snf1p, and based on this, it was expected that mutant *snf4*Δ shows a phenotype similar to that of *snf1*Δ. However, at 1% glucose the mutant *snf4*Δ exhibited higher mitochondrial respiration than the WT strain, and this phenotype is also seen in 10% glucose (Fig. 6). On the contrary, the *GAL83* deletion has no impact on mitochondrial respiration (Fig. 6). Interestingly, deletion of *SNF4* or *GAL83* resulted in increased growth of *S. cerevisiae* even at low glucose concentration (0.1%) (Fig. 7). These data suggest that Snf1p has a specific role in growth and mitochondrial respiration of *S. cerevisiae*, independent of the SNF1 complex.

**Figure 6.**
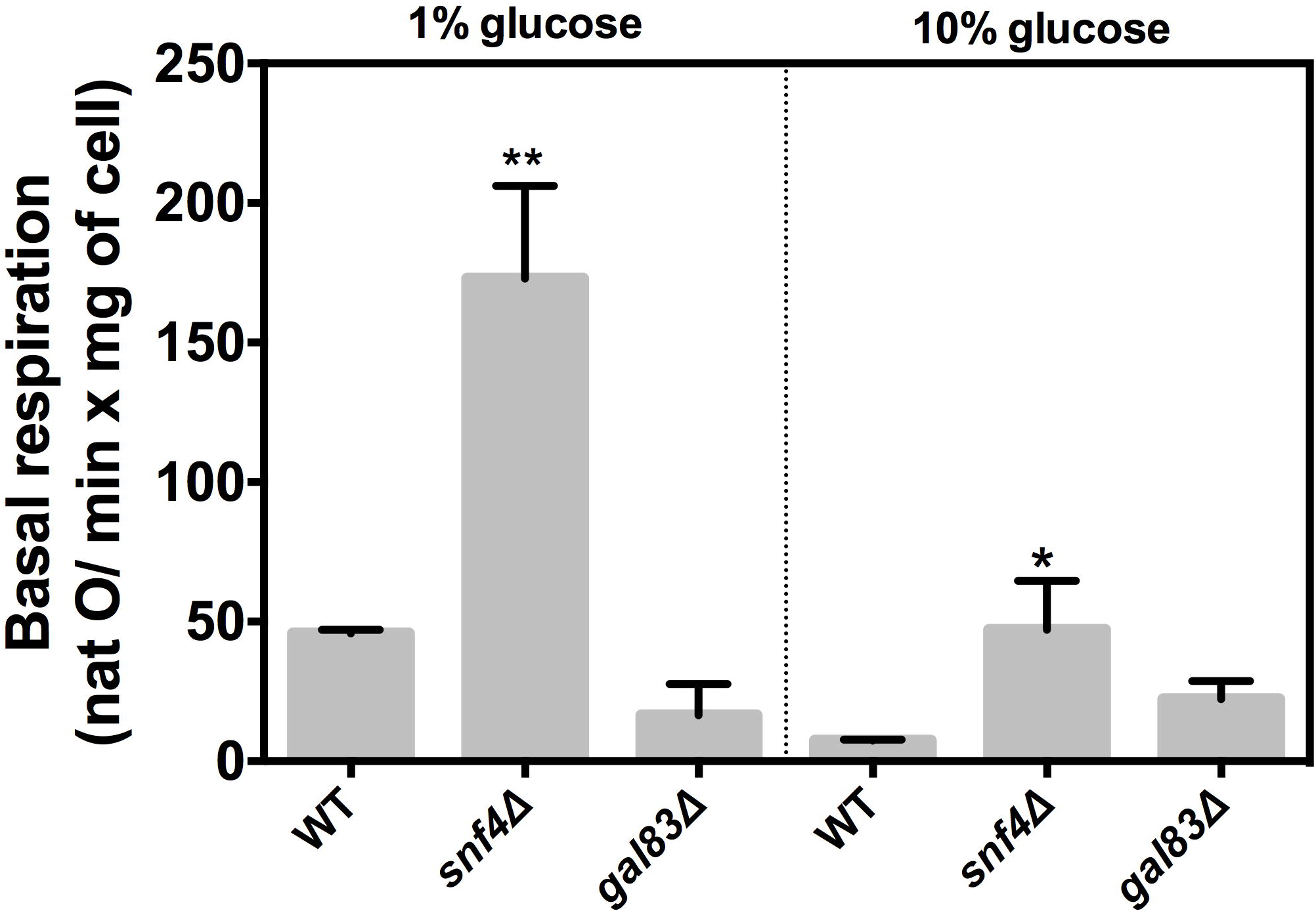
Impact of *SNF4* and *GAL83* deletion in the mitochondrial respiration. The oxygen consumption was measured at basal state and maximal respiratory capacity in exponential phase. The results represent mean values ± SEM from 3-5 independent experiments, which includes mean values of 3 technical repetitions. Statistical analyses were performed using two-tailed unpaired Student *t-*test; ns, no significance.

**Figure 7.**
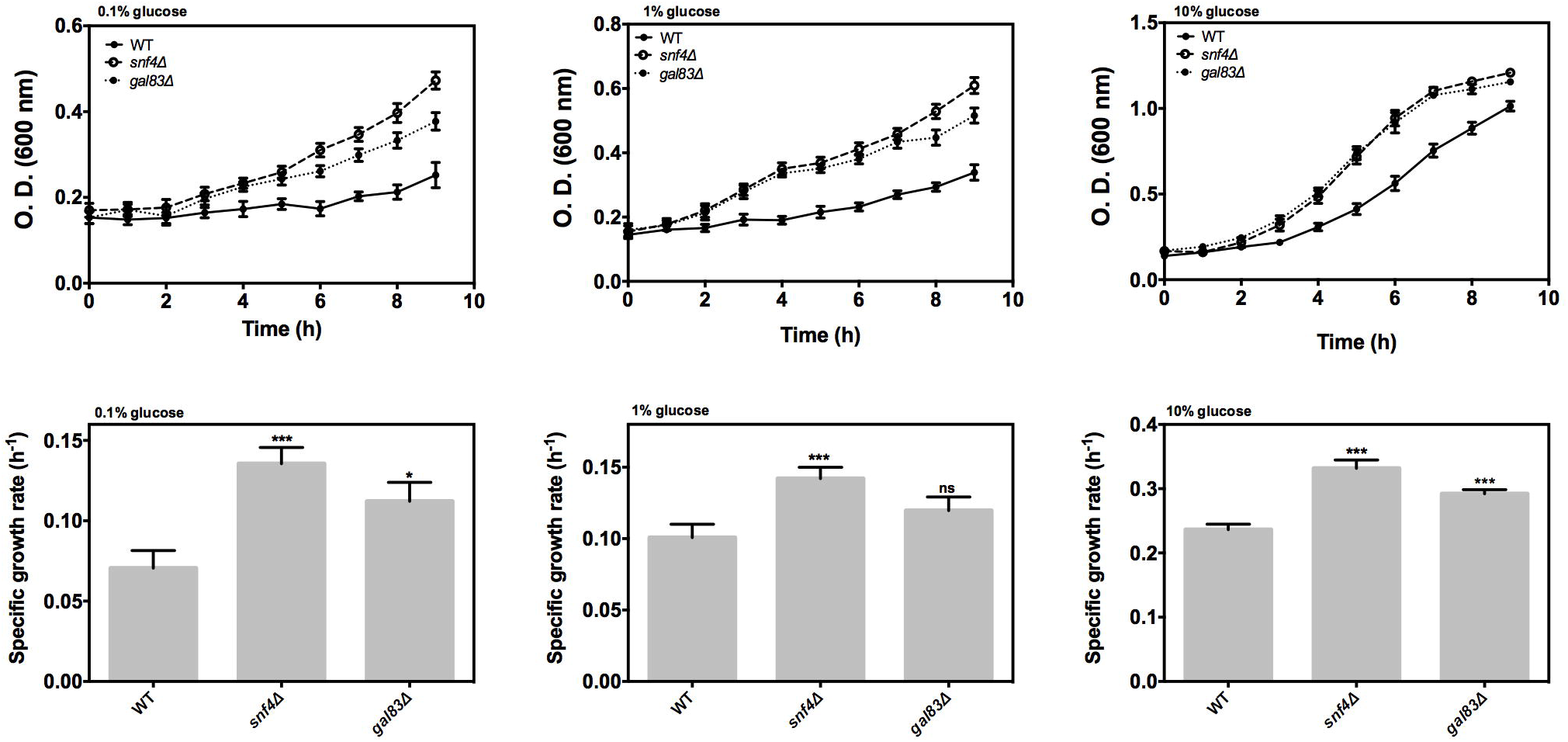
Effect of genes *SNF4* and *GAL83* on cells growth. The growth curve of *S. cerevisiae* growth with 0.1%, 1%, and 10% glucose. The results represent mean values ± SEM from four independent experiments, which includes mean values of three technical repetitions. Statistical significance was calculated by one-way ANOVA followed by Dunnet’s test (\**P* < 0.05 vs. WT; \*\*\**P* < 0.0001 vs. WT; ns=not significant).

## Discussion

The molecular mechanism behind the Crabtree effect remains largely elusive. Control and regulation of this phenotype could help improve ethanol production and establish the basis for the Warburg effect, which is essential in the molecular etiology of cancer. The inhibition of Snf1p by glucose repression indicates that this protein may regulate the Crabtree effect. However, a clear relationship between the Crabtree effect and Snf1p has not been found yet. Cells glucose uptake is the main factor associated with the Crabtree effect and is regulated by hexose transporters mainly, and in turn the Snf1p/Mig1p pathway regulates some hexose transporters transcriptionally. These data imply that Snf1p could regulate the glycolytic flux and the Crabtree effect, and our results support this idea. The *SNF1* mutant is unable to grow in media with low concentration of carbon sources. At 1% glucose, the mutant *snf1*Δ showed a decrease in the glycolytic flux, increased NAD(P)H concentration, enlarged transcription of the *HXK2* gene and reduced mitochondrial respiration. However, the *SNF1* deletion keeps the NADH/NAD^+^ ratio, and the NAD(P)H concentration constant. Also the transcription of genes *HXK2* and *PFK1* at 10% glucose did not show any change, but only the mitochondrial respiration increased. Although the *SNF4* deletion maintains Snf1p inhibited, the phenotypes observed in mitochondrial respiration and growth were different from the *SNF1* mutant, associated to an increase at 1% glucose. In the case of the *GAL83*, we did not detect any change in the mitochondrial respiration or growth. Altogether these findings indicate that *SNF1* is important in the switch of mitochondrial respiration to fermentation.

The signal pathway governed by Snf1p is responsible for the reshaping of the cell metabolism to obtain energy during starvation or in a low-energy environment. In this sense, Snf1p responds primarily to carbon availability and its concentration, irrespective of other carbon or nitrogen sources in the media. It is important to note that concentration of the carbon source is the key factor in *SNF1* mutant growth. This phenotype was also observed in a glucose-repressible carbon source, such as galactose whose canonical regulation pathway is modulated by the Snf1p/Mig1p pathway. This particular phenotype could be explained by the fact that galactose uptake and metabolism are not exclusively regulated by the Snf1p/Mig1p pathway, but also by its proportion relative to glucose (Escalante-Chong, *et al.*, 2015). It has been shown that the extracellular glucose modulates the glucose transporters at a transcription level (Ozcan & Johnston, 1999, Reijenga, *et al.*, 2001). Moreover, the signaling pathway Snf1p/Mig1p repressed high-affinity transporters at high glucose levels. On the contrary, when Snf1p is activated at low glucose concentration, the *HXT6* and *HXT7* genes are de-repressed. Transport of the carbon source is the primary regulator of the glycolytic flux; these data indicate that reduced growth of the *SNF1* mutant could be due to impairment of the glycolytic flux function. Interestingly, the glycolytic flux is only disturbed when the *SNF1* mutant was grown in 1% glucose, and no changes were observed at 10% glucose. This phenotype highlights the importance of *SNF1* at low glucose concentrations. However, it was also reported that at 2% glucose, *SNF1* mutant grown in synthetic medium showed similar glucose consumption as the WT strain (BY4741), but glucose consumption increased at 5 % concentration, relative to the WT strain (Nicastro, *et al.*, 2015). Differences from previous reports could due to the different medium used. Nevertheless, it was observed that *snf1*Δ cells produced fermentation by-products other than ethanol and acetate when grown at 2% glucose and the same amounts were produced in 10% of glucose, when compared to the WT strain (Nicastro, *et al.*, 2015). We did not find any difference in the NADH/NAD^+^ ratio when the WT and *snf1*Δ cells were grown in 1% and 10% glucose. To discard changes in the NADPH species we measured the NAD(P)H fluorescence, detecting an increased concentration in the *snf1*Δ mutant at 1% glucose, whereas no difference was detected at 10% glucose. According to these data NADPH increased in the mutant growing in 1% of glucose. To gain insights into this change the transcription of two key genes in the glycolysis pathway, *HXK2*, and *PFK1* was evaluated. An increase in *HXK2* transcription at 1% glucose in *snf1*Δ cells was the only change found. The increased levels of NADPH and *HXK2* gene transcription suggests that high quantities of glucose-6-phosphate are produced and metabolized via the pentose phosphate pathway. A report supporting this assumption measured the expression of two low-affinity glucose transporters (Hxt1p and the recombinant *TM6*) in a *S. cerevisiae* strain of without hexose transporters. They found increased concentration of glucose-6-phosphate, and fructose-6-phosphate and decreased levels of fructose-1,6-biphosphate, those concentrations might inhibit the activity of the Pfk1p enzyme (Bosch, *et al.*, 2008). Remarkably, the recombinant strains that express the low-affinity glucose transporters experience a lessening in the consumption rate of glucose and fructose, and show a fully respiratory metabolism (Bosch, *et al.*, 2008). In this study we found that mitochondrial respiration decreased in the *snf1*Δ mutant growing in 1% glucose, while at 10% glucose the growth was higher than that exhibited by the WT. The increased mitochondrial respiration at 10% glucose clearly showed the importance of *SNF1* in the switch of mitochondrial respiration to fermentation to obtain ATP. At 10% glucose Snf1p is inhibited by the glucose repression, so we expected that at this concentration the *SNF1* deletion showed no metabolic effect according to all parameters assayed. However, the fact that *SNF1* mutation increased mitochondrial respiration even at high glucose concentration suggests the important function of this gene even in high concentrations of this carbohydrate. In this regard, it has been proposed that at high glucose concentrations Snf1p could form complexes, which in turn would have activated a different set of transcriptional factors implicated in the adaptation of cells growth in fermentation conditions. This phenotype is evident in the transcriptional regulation of the *TPK1* gene by the Snf1p/Cat8p pathway (Galello, *et al.*, 2017). Another example of this model is the endocytosis of the high-affinity glucose transporter Hxt6p at high glucose concentrations that requires the arresting-related trafficking adaptor Rod1p/Art4p, whose function is regulated by Snf1p (Llopis-Torregrosa, *et al.*, 2016). On the contrary, the decrease in mitochondrial respiration observed at 1% glucose could be related to reduction of the glycolytic flux and cellular viability in the mutant *snf1*Δ. Altogether, these results indicate that *SNF1* gene is important to modulate the glycolytic flux and mitochondrial respiration, which is essential at low carbon sources concentration, whereas at higher glucose concentrations *SNF1* gene is critical for the establishment of fermentation.

Finally, these findings suggest that *SNF1* controls the glycolytic flux and mitochondrial respiration through regulation of the hexose transporters, which in turn allow switching from mitochondrial respiration to fermentation.

## Acknowledgments

The authors would like to thank Minerva Ramos-Gomez for their technical support. The authors declare no competing financial interest.

